# Identifying site- and stimulation-specific TMS-evoked EEG potentials using a quantitative cosine similarity metric

**DOI:** 10.1101/612499

**Authors:** Michael Freedberg, Jack A. Reeves, Sara J. Hussain, Kareem A. Zaghloul, Eric M. Wassermann

## Abstract

The ability to interpret transcanial magnetic stimulation (TMS) evoked EEG potentials (TEPs) is limited by artifacts, such as auditory evoked responses produced by the TMS coil. TEPs generated from direct cortical stimulation should vary in spatial distribution with stimulation site and from responses to sham stimulation. Responses that do not show these effects are likely to be artifactual. In 20 healthy volunteers, we delivered active and sham TMS to right prefrontal, left primary motor, and left posterior parietal cortex and compared the waveform similarity of TEPs between stimulation sites and active and sham TMS with a cosine similarity-based analysis method. We looked for epochs after the stimulus when the spatial pattern of TMS-evoked activation had greater than random similarity between stimulation sites and sham vs. active TMS, indicating presence of a dominant artifact, such as the auditory brain response. We calculated and binarized the derivatives of the TEPs recorded from 30 EEG channels and calculated cosine similarity between conditions at each time point with millisecond resolution. Only TEP components occurring before approximately 80 ms differed across stimulation sites and between active and sham, indicating site and condition-specific responses. TEP components before about 80 ms can be safely interpreted as stimulation location-specific responses to TMS, but components beyond this latency should be interpreted with caution due to high similarity in their spatial distribution.

## Introduction

Transcranial magnetic stimulation (TMS)-evoked EEG potentials (TEPs) are a rich source of neurophysiological data, and can be used to study cortical excitability and effective connectivity [1]. TEPs are obtained by activating the cortex with TMS and measuring the response with electroencephalography (EEG) across the scalp. TEPs have been used to study a variety of phenomena, including consciousness [2,3], memory [4], and the pathophysiology of neurological disorders [5–7].

A major factor limiting the interpretation of the EEG response to TMS is the presence of time-locked artifacts. Artifacts from sources outside of the brain, such as electrode noise and recharge artifacts from the TMS coil, do not interact directly with brain signals and can be removed offline with techniques such as independent component analysis (ICA) [8,9]. However, artifacts from cortical responses to auditory or somatosensory stimulation by the TMS pulse can interact with and modify the amplitude of other neural signals [10,11]. Therefore, sensory artifacts cannot be isolated and removed reliably with ICA [12] or by subtracting the amplitude of sham-evoked auditory and somatosensory artifacts from active TMS responses. Sensory artifacts can be attenuated with noise masking and vibration-damping of the coil [2,13]. However, a recent study [14] showed that data collected under these conditions may still be contaminated by artifact.

Without the ability to remove brain-generated artifacts, such as auditory and somatosensory responses, it is important to identify epochs of the TMS response which contain these artifacts and interpret signals from those epochs with caution or avoid them altogether. Previous studies have shown that later TEP components, e.g., the N100, do not differ significantly in their scalp distribution by stimulation site or between sham and active stimulation [15,16]. This lack of spatial selectivity is inconsistent with a response of the brain to focal stimulation [16]. It is likely, therefore, that N100 and later components are contaminated by sensory artifact or some other non-selective process [13,17,18]. A study using noise masking and reduction of sensation with foam material between the coil and scalp [16] supported this idea by showing significant correlations between active and sham-evoked waveforms at latencies corresponding to the N100 component. Correlated waveforms indicate components that are common across stimulation conditions (e.g. active and sham) in amplitude-independent timing and pattern of evoked activity. Correlation analysis has an advantage over amplitude-based methods, which only compare the voltage at a single electrode between conditions [16] and can lead to misidentification of waveforms as local brain responses, due to ill-controlled or incompletely removed artifact. However, correlation analysis has lower time resolution. These issues can be avoided by calculating the instantaneous cosine similarity between waveforms. Cosine similarity compares multiple channels and can be used to measure similarity in the shapes of responses across the scalp (between-condition similarity). The cosine method can also be used to measure the consistency of responses within stimulation sites and conditions (repeatability).

In this study, we used cosine similarity to quantify between-condition similarity and repeatability of responses to active and sham TMS delivered to three cortical sites. We did not apply noise masking during data collection, allowing us to determine whether site-specific signal can be separately identified from responses to the auditory stimulus from the TMS coil. Our goal was to identify epochs were the signal was differentiated between stimulation conditions and locations, indicating that the response was caused by cortical activation at the stimulation site.

## Methods

### Participants

Twenty healthy individuals (11 F; mean age 24.3 ± 3.1 years) gave informed consent to participate in the study, which was approved by the NIH Combined Central Nervous System Institutional Review Board. All experiments were performed in accordance with the Declaration of Helsinki.

### MRI acquisition and TMS targeting

MPRAGE T_1_-weighted scans were obtained from each participant using either a GE 3T HDx scanner (N = 16, 32-channel head coil, voxel size = 1 cm^3^, field of view = 25.6 cm^2^, flip angle = 7°, 172 transverse slices) or a Siemens 3T Skyra scanner (N = 4, 32-channel head coil, voxel size = 0.926 cm x 0.926 cm x 0.900 cm, field of view = 23.7 cm^2^, flip angle = 9°, 192 sagittal slices). Predetermined TMS target locations for the right dorsolateral prefrontal cortex (DLPFC; MNI: 42 46 4) [19] and left posterior parietal area (PPC; MNI −48 −65 45) [20] were prepared for each participant by transforming their anatomical scans to MNI space, placing a 3 mm radius circular marker at the target location, and transforming the scans back to subject space using AFNI neuroimaging analysis software [21]. The target location for the primary motor area (M1) was determined empirically (see “TMS and EEG recording” section). For PPC stimulation, the handle of the coil was rotated approximately 10° laterally from the antero-posterior axis [20]. For DLPFC stimulation, the handle of the coil was angled 45° medially from the anterior-to-posterior axis. For M1 stimulation, the coil angle was adjusted to the orientation that maximized the motor response (see “TMS and EEG recording” section).

### TMS, EMG, and EEG recording

A Brainsight frameless neuronavigation system was used to guide stimulation to the left PPC, left M1, and right DLPFC targets in each participant. Monophasic TMS was delivered with a 70 mm figure-of-eight coil connected to a Magstim 200 stimulator. Electromyographic responses were measured from the right first dorsal interosseous muscle with standard surface electrodes. The reference electrode was placed over the metacarpophalangeal joint of the index finger. The target for M1 stimulation was the scalp site where stimulation required the lowest intensity to evoke at least five motor-evoked potentials (MEPs) > 50 uV on ten consecutive trials. Resting MEP threshold (RMT) was calculated using an automatic threshold-tracking algorithm (TMS Motor Threshold Assessment Tool, MTAT 2.0) [22]. Stimulation intensity for active and sham stimulation at all sites was set to each participant’s RMT.

### Experimental design and recording

We delivered 150 active and 50 sham TMS pulses at each stimulation site. The order of sites was counterbalanced between participants. Pulses were delivered every five seconds with a 15% jitter to reduce any effects of pulse anticipation. The EEG was sampled at 5000 Hz (DC filtered at 1000 Hz) from 30 recording scalp electrodes in a 10-20 arrangement using a TMS-compatible EEG system (BrainAmp MR+, Brain Vision). Electrode impedances were kept < 10 kΩ. The goal of sham stimulation was to reduce cortical stimulation *without eliminating the auditory artifact*. This was achieved by placing a 3 cm cardboard spacer between the coil and scalp. For each location, active TMS was performed first, followed by sham. Participants wore foam earplugs during all TMS procedures.

### Data preprocessing

We preprocessed the data with a standard analysis pipeline (TMS-EEG Signal Analyzer (TESA) extension for EEGLAB [8] running in a MATLAB environment). Trials were epoched from −1.5 to +3 sec around each TMS pulse and data from −1 ms to +15 ms around each pulse were removed and replaced with a cubic interpolation. The data were down-sampled to 1000 Hz and noisy channels were automatically detected and removed if they exceeded a full-epoch kurtosis measurement of 5. On average, we removed one channel per participant for each stimulation condition (minimum = 0; maximum = 3). We baseline-corrected each epoch by whole-epoch de-meaning and then performed two rounds of ICA to detect and remove artifacts. Both rounds used the FastICA algorithm [23] with a symmetric decomposition approach and hyperbolic tangent contrast function. The first round of ICA included an automated artifact detection algorithm (tesa_compselect) [8] to remove large pulse-related artifacts. The second round was performed to eliminate small artifacts not rejected during the first round. To obtain a consensus about which components should be rejected, authors JR, SH, and MF performed agreement training on a test data set. Each author individually performed components rejection on subsets of test data until substantial inter-rater agreement was achieved (> 0.78 Cohen’s kappa interrater coefficient) [24,25]. Author JR performed the final component rejection for the second round of ICA. Between the two rounds of ICA, we processed the data with a 1-100 Hz zero-phase bandpass Butterworth filter and a 58-62 Hz zero-phase band-stop filter to remove line noise. Following ICA, we baseline-corrected individual trials from −100 ms to −10 ms relative to TMS pulse. Channels were re-referenced to the average of all channels for each trial. Finally, we calculated the time-locked average waveforms for each participant and each condition and visually inspected each participant’s preprocessed TEPs by plotting the trial-average for each EEG channel. We performed an additional data quality check by inspecting plots of group-level channel averages (Fig 1).

**Fig 1.**
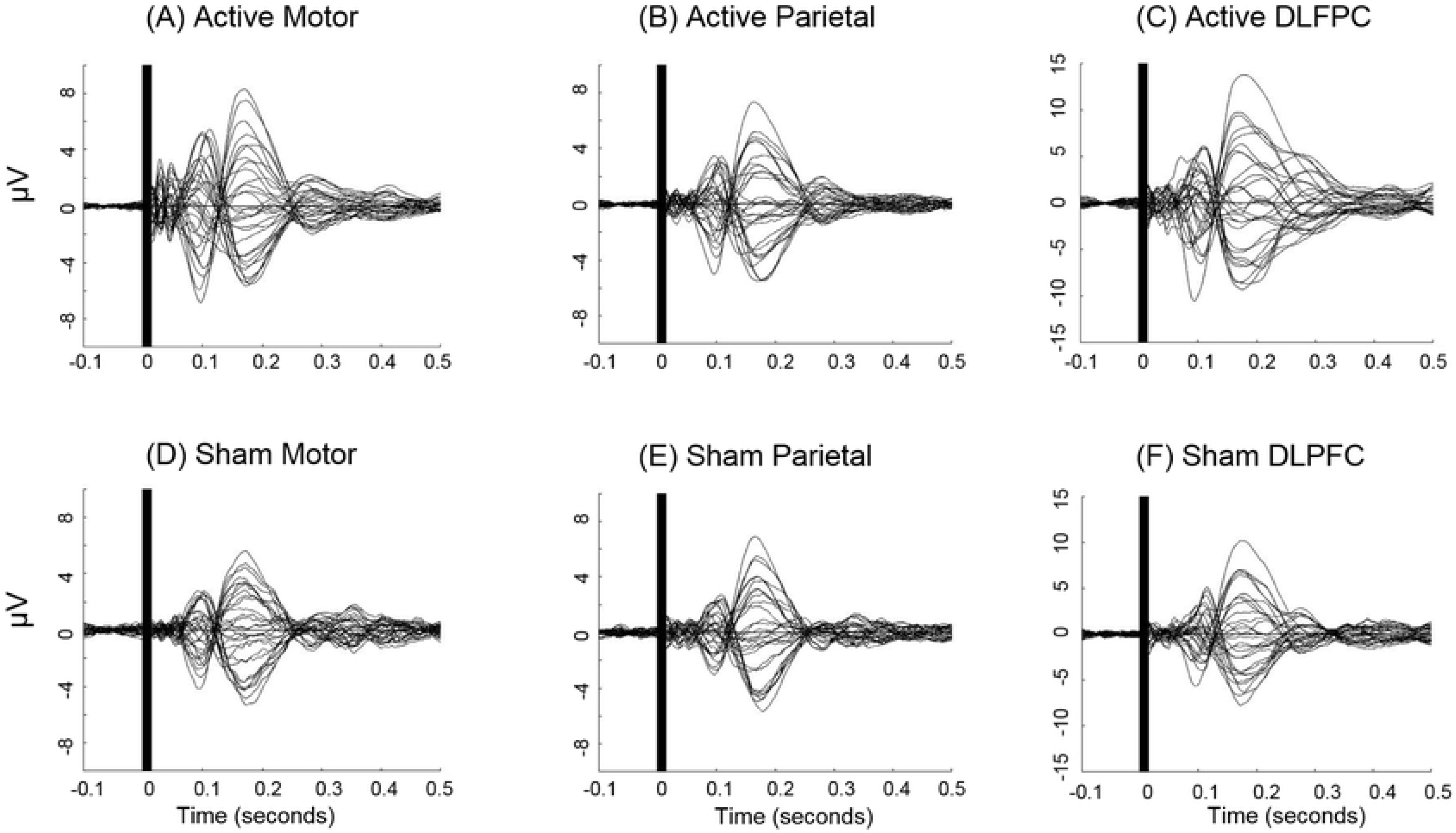
TEPs for each of the six stimulation conditions. Group-averaged EEG voltage traces for active and sham M1 (A and D), PPC (B and E), and DLPFC (C and F) stimulation. Plots show average voltage traces for EEG channels FP1, FP2, F3, F4, C3, C4, P3, P4, O1, O2, F7, F8, T7, T8, P7, P8, FZ, CZ, PZ, IZ, FC1, FC2, CP1, CP2, FC5, FC6, CP5, CP6, TP9, and TP10.

### Similarity analysis

We used a modified cosine similarity method [26] to measure within- and between-condition similarity in topographical activity patterns. We used a within-subjects design to compare evoked responses for each participant separately, with statistically significant differences calculated at the group level (see “Statistical analyses of similarity measurements,” below). To identify similar patterns, we compared TEPs between active to sham stimulation conditions at each stimulation site and active to active stimulation between stimulation sites, resulting in six sets of similarity measurements. We also computed the repeatability of responses to active stimulation at each site by splitting the trials within the same stimulation condition into two samples and comparing them using the cosine method. Repeatability analysis provided a practical maximum level of similarity for that site to contrast with the similarity in the between-condition and location comparisons. This accounted for noise and other factors which caused the maximum to decrease from the theoretical value of 1.

For each condition, we created an average waveform from 50 randomly-selected trials from each participant. We used 50 trials to equalize the number of trials between comparisons (active vs. sham and between-location active stimulation), since only 50 sham trials were delivered. Non-overlapping trials were used for within-condition comparisons. We took the derivatives of the preprocessed and averaged TEPs at every time point for each channel by subtracting the amplitude of each timepoint from the amplitude of preceding one (Fig 2A). The derivatives were then binarized so that an increase in point-to-point amplitude was assigned a value of +1 and a decrease a value of −1 (Fig 2B). This removed the amplitude dimension of the data and allowed for model-free analysis of the shape of the waveform. For each comparison, cosine similarity between the two “feature vectors” [26] was calculated (Fig 2C) at each time-point using all channels as follows:

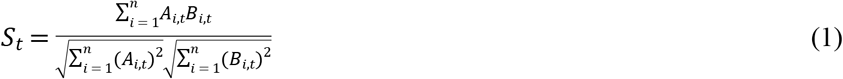

where *S*_*t*_ is cosine similarity at time *t*, *A*_*i,t*_ and *B*_*i,t*_ are the binarized derivatives for each condition at channel *i* and time *t*, and *n* is the total number of channels. The process was repeated 1000 times for each participant and each comparison, using randomly-permuted trial averages. The within-participant mean of the averages was then calculated as the measure of similarity between active and sham conditions and between stimulation sites (“between-condition similarity”), and TEP repeatability.

**Fig 2.**
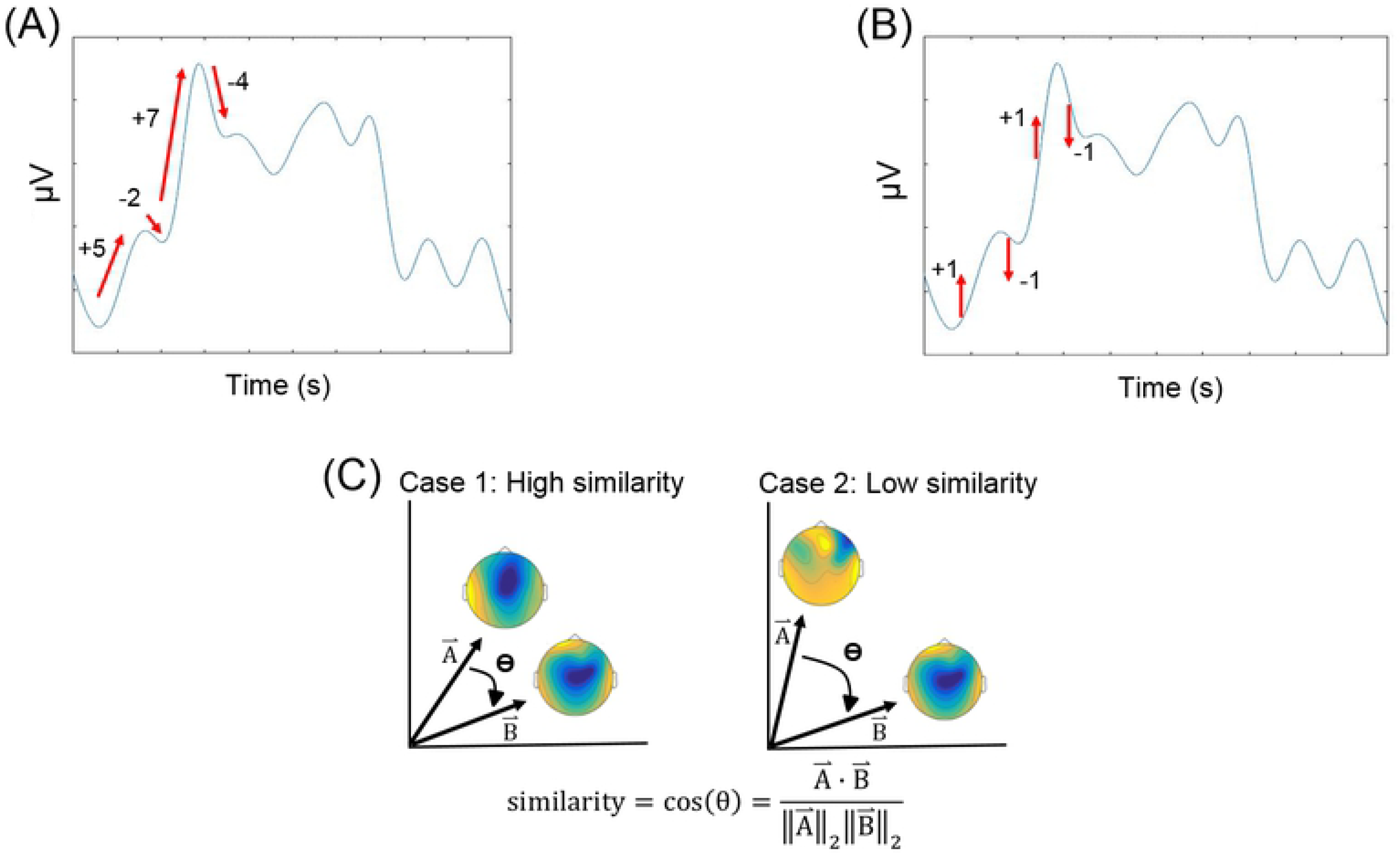
Visualization of cosine similarity method. We took the derivatives of the preprocessed and averaged TEPs at every time point for each channel (A), binarized the derivatives so that an increase in point-to-point amplitude was assigned a value of +1 and a decrease a value of −1 (B), and calculated cosine similarity between the two “feature vectors” (C).

### Statistical analyses of similarity measurements

For the similarity analysis (see above), we used cluster-based permutations [27] to perform group-level comparisons within and between conditions. Two types of tests were conducted: comparisons of each similarity curve to pre-stimulus baseline values and comparisons of within-condition and between-condition similarity curves.

We performed statistical comparisons to reveal significant differences between baseline and post-TMS repeatability, and between-condition similarity. For each within- and between-condition comparison, each participant’s average similarity curve was segmented into baseline (1500—500 ms before TMS) and response (15—1015 ms after TMS) epochs. A *p-*value was calculated for each time-point by comparing the distributions of the participants’ baseline and response epochs. Clusters of significant time-points were identified by summing the *p-*values of all contiguous time-points that reached a threshold of 0.05. We controlled for multiple comparisons by comparing each of the cluster sizes to a distribution of 1000 *p-*value clusters derived from randomly-permuted epoch comparisons and retaining only original clusters > 95% of the random-cluster distribution. Each random cluster was created by comparing two randomly-sampled distributions of twenty epochs each from the combined pool of the baseline and response epochs and selecting the size of the largest cluster (by summed *p-*value, as above).

## Results

### Group-level TEPs

Active TMS produced significant cortical activation at the M1, PPC, and DLPFC sites, with maximum group-level peak-to-peak amplitudes across the scalp of 13.9 μV, 12.9 μV, and 23.0 μV, respectively (Fig 1). Sham TMS also produced significant activation at all three sites but with lower amplitude than active stimulation (M1: 10.9 μV, PPC: 12.5 μV, DLPFC: 17.9 μV).

Qualitatively, the TEPs were spatially differentiated between 15 ms (end of interpolation) and 60 ms (Fig 3). Between 60 and 80 ms, the patterns started to become similar between conditions. By 100 ms, the responses became highly similar and remained so until approximately 300 ms.

**Fig 3.**
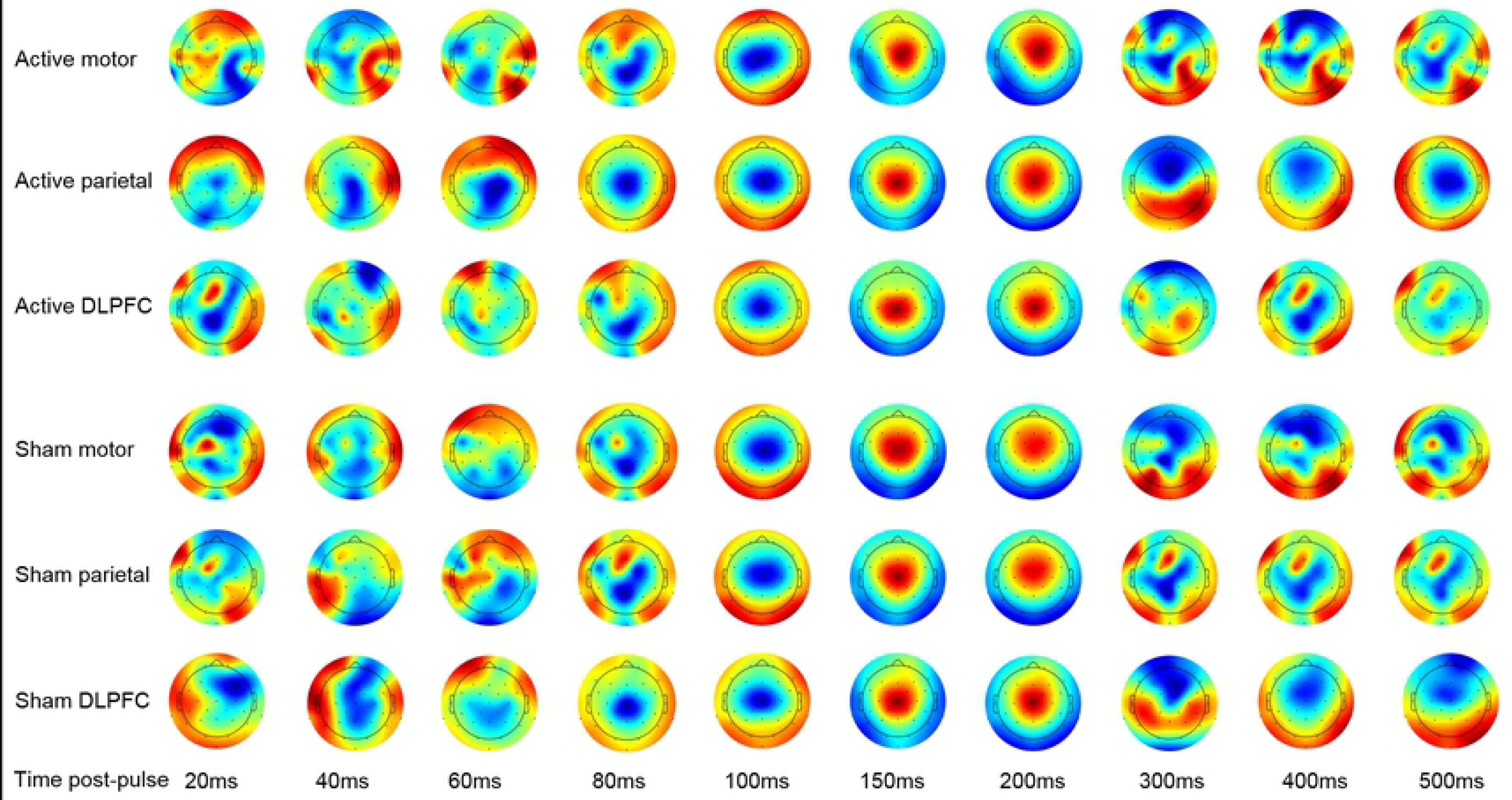
Topographic patterns of activation for all six conditions. Group-level topographic activity plots for each of the six stimulation conditions (active M1, active PPC, active DLPFC, sham M1, sham PPC, and sham DLPFC). Each plot is scaled to the minimum and maximum electrode voltage at that timepoint to emphasize the patterns of activation.

### Repeatability of topographical patterns of activation

The repeatability of TEP waveforms was maximal immediately following the pulse for all stimulation sites and decayed to pre-stimulus values by 500 ms after the pulse (Fig 4). Significant differences from baseline started at 15 ms (the end of interpolation) and extended to 313 ms for M1, 289 ms for PPC, and 384 ms for DLPFC active stimulation. A large drop in repeatability occurred at a mean of 136 ms after TMS for all three locations.

**Fig 4.**
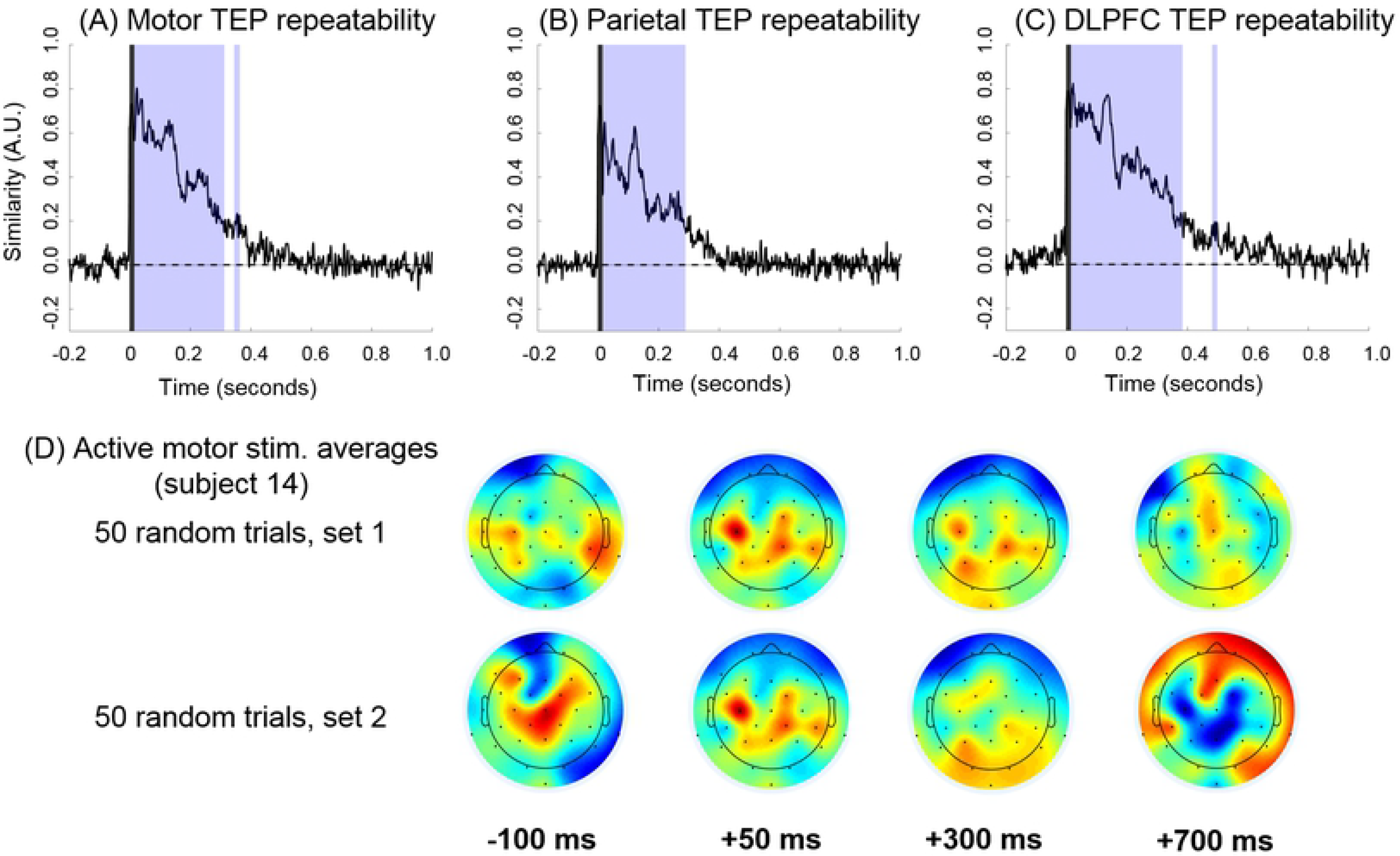
Temporal and spatial results of repeatability analyses reflecting similarity within active stimulation conditions. (Top) Group-level repeatability curves comparing within-participant and within-condition averages for active M1 (A), PPC (B), and DLPFC stimulation (C). The y-axis indicates arbitrary similarity units with a possible range of −1 to +1, with +1 indicating perfect similarity, −1 indicating perfect anti-similarity, and 0 indicating complete non-similarity. The lightly shaded regions correspond to significant differences from pre-stimulation baseline across time, indicating topographic patterns that are nonrandom. The darkly shaded regions from −1 ms to 15 ms indicate TMS pulse interpolation. (Bottom) Topographical distributions of two active-M1 averages (constructed using 50 non-overlapping trials each) from a representative participant (D).

### Between-condition similarity

Two increases in similarity appeared in every between-condition (active vs sham, between-location) comparison (Fig 5). The first started at a mean of 84 ± 13 ms after the TMS pulse and peaked at 128 ± 7 ms. The second started at 172 ± 7 ms and peaked at 229 ± 16 ms. Significant similarity was present prior to 84 ± 13 ms post-pulse in all but the M1-DLPFC active-comparison. However, unlike the later epochs, the specific epochs showing similarity before 84 ms were different between comparisons, indicating site specific patterns of activity rather than stereotyped signal from artifact. Overall, similarity between all conditions decayed to baseline values by 350 ms (Fig 5).

**Fig 5.**
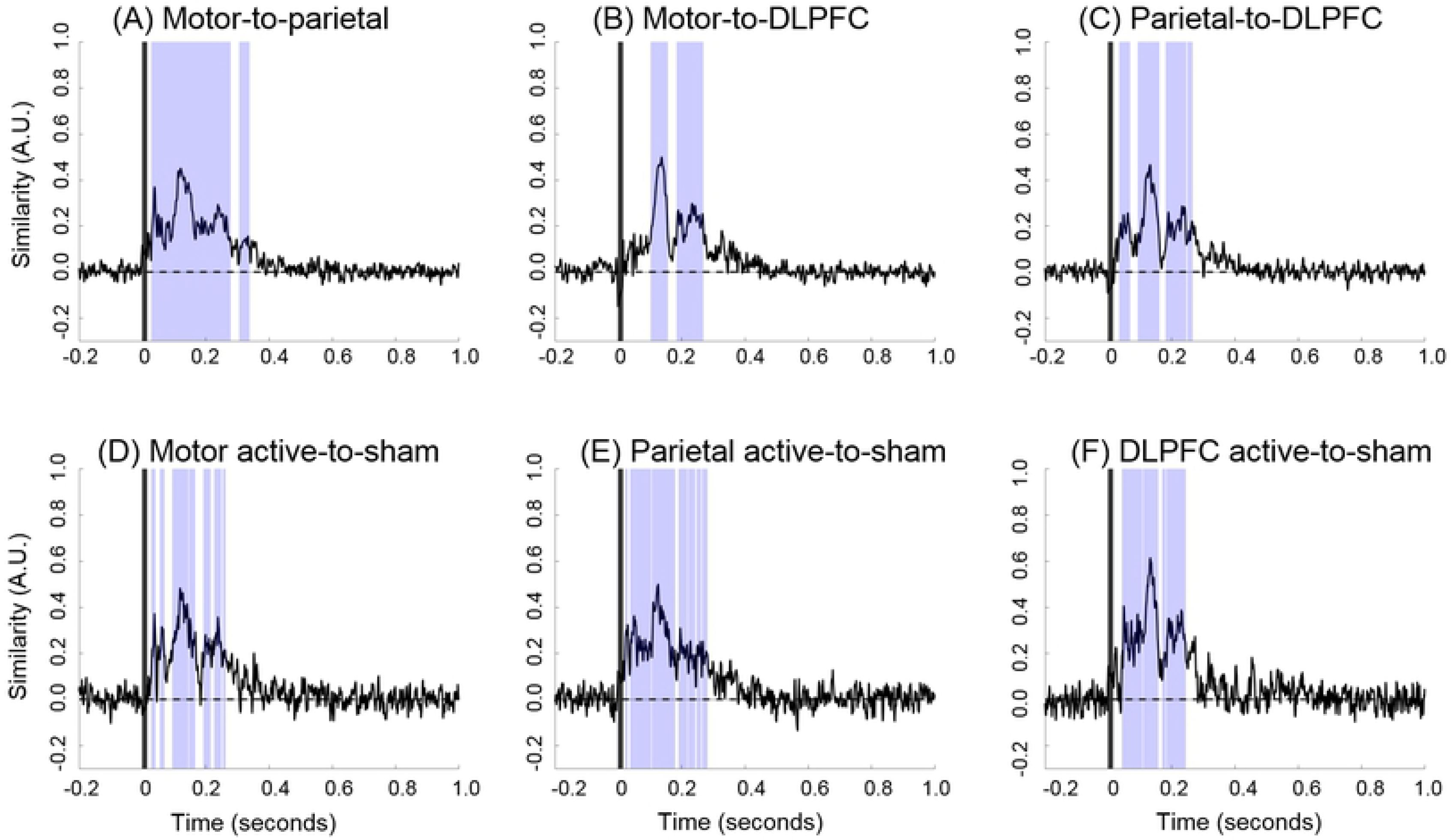
Between-condition similarity curves. Between-location, active-to-active similarity comparisons for M1-PPC (A), M1-DLPFC (B), and PPC-DLPFC (C) and active-sham similarity comparisons at M1 (D), PPC (E), and DLPFC (F) locations. Light gray shading reflects regions of significant difference relative to baseline, indicating the similarity between the compared conditions is above chance. The darkly shaded regions from −1 ms to 15 ms indicate TMS pulse interpolation.

## Discussion

We measured similarity in the spatial distributions of TEPs from active and sham TMS and from TEPs from active stimulation at different sites using the cosine similarity method. Comparisons across these conditions allowed us to identify epochs at which TEP patterns of activation exhibited above-chance similarity. Greater-than random similarity between responses to active and sham stimulation indicates the presence of artifact, probably from auditory and/or somatosensory stimulation by the TMS pulse. From our between-condition similarity analysis, we obtained results consistent with the previous finding [14] that early responses (20-80 ms after TMS) are significantly correlated in waveform shape between stimulation conditions (Fig 5). However, in our results, the specific 20-80 ms epochs with statistically significant similarity differed between comparisons (stimulation site, active vs. sham). For example, active M1 was significantly similar to active PPC from 28-279 ms after TMS, while active M1 was not significantly similar to active DLPFC until 102 ms after TMS. This indicates that the events in the period from 20-80 ms did not contain an artifact common to all stimulation conditions (i.e., auditory artifact), which is consistent with claims [16,17] that TEP components occuring 20-80 ms after the TMS pulse are condition and site specific. We note, however, that somatosensory effects are likely to vary by stimulation site and cannot be distinguished from signal caused by direct cortical stimulation. Across all comparisons, there was a rapid change from low to high similarity at 84 ± 13 ms, corresponding with the onset of the N100 component. Our findings comport with the concept of the N100 as a nonspecific response to cortical stimulation [14,15,28,29].

We found decreasing repeatability of whole-brain TEP patterns with time after the TMS pulse (Fig 4), indicating that within-session latency and waveform shape reproducibility of the TEP decays with time after the TMS pulse. We observed a large drop in repeatability at an average of 136 ms post-pulse, indicating that the latencies of TEP components occurring before 136 ms can be precisely estimated in our data, while later latencies may require higher number of trials to estimate with similar precision. These results are consistent with those of a previous study [30], which showed higher between-session repeatability in peak amplitude and latency for early components (≤ 100 ms) than for a late component corresponding to the P180. However, our results differ with those of a later study [31] which found P180 amplitude to be equally repeatable as the early N40 and P60 components for within-session comparisons. A possible reason for this discrepancy is that the latter study [31] analyzed a subset of electrodes which was biased toward the vertex and prone to sensory contamation [28], which could artificially raise the repeatability of the response. It is important to note that these studies used amplitude-dependent repeatability measures. Our cosine-based repeatability results quantify the consistency in the timing and response direction of the spread of evoked activity, factors which are not explicitly accounted for in amplitude-based studies.

The objective of this study was to use cosine similarity to distinguish between responses to cortical stimulation and neurally generated sensory artifacts in the TEP, therefore we did not attempt to mask the auditory stimulus. This means that any difference in the TEPs between active stimulation locations, and between active and sham stimulation, was due either to direct cortical stimulation or differences in somatosensory stimulation. Even though noise masking reduces the amplitude of TEPs, particularly the N100 component, it does not alter the fundamental shape of the waveform [13,17], possibly due to incomplete masking of the TMS acoustic stimulus [14]. Cosine similarity could be used as a test for the effectiveness of sensory masking.

There are reasons to not apply noise masking in TEP studies. Background noise can have differential effects on task performance between experimental groups, such as children diagnosed with ADHD vs healthy controls [32], introverts vs extroverts [33], and can affect performance on cognitive vs. perceptual tasks differently [34]. For studies that do not use active sound masking methods [4,35], our results indicate that early-latency (<80 ms) TEPs are reliably location specific.

## Conclusions

Although artifacts may be present over the entire TEP response, early TEP components best reflect site and condition-specific responses to cortical stimulation. However, later responses become progressively similar between stimulation conditions, suggesting that these components predominately reflect neural artifacts. We propose that unless it can be shown that sensory masking eliminates residual neurally generated artifact, only TEP components occurring earlier are safely interpretable as stimulation location-specific responses to TMS cortical stimulation. In addition, the reproducibility of the TEP waveform decays monotonically from the TMS pulse. While later components may contain valuable physiological information, we suggest caution in in their interpretation and warn against their attribution to direct cortical stimulation by the TMS pulse. Auditory and somatosensory stimulation controls and/or further refinements in signal processing may be necessary to extract true late responses to cortical stimulation.

## References

1. Ilmoniemi RJ, Kičić D. Methodology for combined TMS and EEG. Brain Topogr. 2010;22: 233–248. doi:10.1007/s10548-009-0123-4

2. Massimini M, Ferrarelli F, Huber R, Esser SK, Singh H, Tononi G. Breakdown of cortical effective connectivity during sleep. Science. 2005;309: 2228–2232. doi:10.1126/science.1117256

3. Ferrarelli F, Massimini M, Sarasso S, Casali A, Riedner BA, Angelini G, et al. Breakdown in cortical effective connectivity during midazolam-induced loss of consciousness. Proc Natl Acad Sci. 2010;107: 2681–2686. doi:10.1073/pnas.0913008107

4. Rogasch NC, Daskalakis ZJ, Fitzgerald PB. Cortical inhibition of distinct mechanisms in the dorsolateral prefrontal cortex is related to working memory performance: A TMS-EEG study. Cortex. 2015;64: 68–77. doi:10.1016/j.cortex.2014.10.003

5. Bruckmann S, Hauk D, Roessner V, Resch F, Freitag CM, Kammer T, et al. Cortical inhibition in attention deficit hyperactivity disorder: New insights from the electroencephalographic response to transcranial magnetic stimulation. Brain. 2012;135: 2215–2230. doi:10.1093/brain/aws071

6. Ferreri F, Vecchio F, Vollero L, Guerra A, Petrichella S, Ponzo D, et al. Sensorimotor cortex excitability and connectivity in Alzheimer’s disease: A TMS-EEG co-registration study. Hum Brain Mapp. 2016;37: 2083–2096. doi:10.1002/hbm.23158

7. Frantseva M, Cui J, Farzan F, Chinta L V., Perez Velazquez JL, Daskalakis ZJ. Disrupted cortical conductivity in schizophrenia: TMS-EEG study. Cereb Cortex. 2014;24: 211–221. doi:10.1093/cercor/bhs304

8. Rogasch NC, Sullivan C, Thomson RH, Rose NS, Bailey NW, Fitzgerald PB, et al. Analysing concurrent transcranial magnetic stimulation and electroencephalographic data: A review and introduction to the open-source TESA software. Neuroimage. 2017;147: 934–951. doi:10.1016/j.neuroimage.2016.10.031

9. Rogasch NC, Thomson RH, Farzan F, Fitzgibbon BM, Bailey NW, Hernandez-Pavon JC, et al. Removing artefacts from TMS-EEG recordings using independent component analysis: Importance for assessing prefrontal and motor cortex network properties. Neuroimage. 2014;101: 425–439. doi:10.1016/j.neuroimage.2014.07.037

10. Foxe JJ, Wylie GR, Martinez A, Schroeder CE, Javitt DC, Guilfoyle D, et al. Auditory-somatosensory multisensory processing in auditory association cortex: An fMRI study. J Neurophysiol. 2002;88: 540–543. doi:10.1152/jn.00694.2001

11. Molholm S, Ritter W, Murray MM, Javitt DC, Schroeder CE, Foxe JJ. Multisensory auditory-visual interactions during early sensory processing in humans: A high-density electrical mapping study. Cogn Brain Res. 2002;14: 115–128. doi:10.1016/S0926-6410(02)00066-6

12. Stam CJ. Nonlinear brain dynamics [Internet]. Nova Biomedical. 2006. Available: http://www.loc.gov/catdir/toc/ecip065/2005037344.html

13. ter Braack EM, de Vos CC, van Putten MJ. Masking the auditory evoked potential in TMS-EEG: A comparison of various methods. Brain Topogr. 2015;28: 520–528. doi:10.1007/s10548-013-0312-z

14. Conde V, Tomasevic L, Akopian I, Stanek K, Saturnino GB, Thielscher A, et al. The non-transcranial TMS-evoked potential is an inherent source of ambiguity in TMS-EEG studies. Neuroimage. 2019;185: 300–312. doi:10.1016/j.neuroimage.2018.10.052

15. Du X, Choa F Sen, Summerfelt A, Rowland LM, Chiappelli J, Kochunov P, et al. N100 as a generic cortical electrophysiological marker based on decomposition of TMS-evoked potentials across five anatomic locations. Exp Brain Res. 2017;235: 69–81. doi:10.1007/s00221-016-4773-7

16. Harquel S, Bacle T, Beynel L, Marendaz C, Chauvin A, David O. Mapping dynamical properties of cortical microcircuits using robotized TMS and EEG: Towards functional cytoarchitectonics. Neuroimage. 2016;135: 115–124. doi:10.1016/j.neuroimage.2016.05.009

17. Nikouline V, Ruohonen J, Ilmoniemi RJ. The role of the coil click in TMS assessed with simultaneous EEG. Clin Neurophysiol. 1999;110: 1325–1328. doi:10.1016/S1388-2457(99)00070-X

18. Herring JD, Thut G, Jensen O, Bergmann TO. Attention modulates TMS-Locked alpha oscillations in the visual cortex. J Neurosci. 2015;35: 14435–14447. doi:10.1523/JNEUROSCI.1833-15.2015

19. Poldrack RA, Clark J, Paré-Blagoev EJ, Shohamy D, Creso Moyano J, Myers C, et al. Interactive memory systems in the human brain. Nature. 2001;414: 546–550. doi:10.1038/35107080

20. Koch G, Ponzo V, Di Lorenzo F, Caltagirone C, Veniero D. Hebbian and Anti-Hebbian Spike-Timing-Dependent Plasticity of Human Cortico-Cortical Connections. J Neurosci. 2013;33: 9725–9733. doi:10.1523/JNEUROSCI.4988-12.2013

21. Cox RW. AFNI: Software for analysis and visualization of functional magnetic resonance neuroimages. Comput Biomed Res. 1996;29: 162–173. doi:10.1006/cbmr.1996.0014

22. Awiszus F, Borckardt J. TMS Motor Threshold Assessment Tool (MTAT 2.0) [Internet]. 2011. Available: http://www.clinicalresearcher.org/software.htm

23. Hyvärinen A, Oja E. Independent component analysis: Algorithms and applications. Neural Networks. 2000;13: 411–430. doi:10.1016/S0893-6080(00)00026-5

24. Hallgren KA. Computing inter-rater reliability for observational Data: An overview and tutorial. Tutor Quant Methods Psychol. 2012;8: 23–34. doi:10.20982/tqmp.08.1.p023

25. McHugh ML. Interrater reliability: The kappa statistic. Biochem Medica. 2012; 276–282. doi:10.11613/BM.2012.031

26. Yaffe RB, Kerr MSD, Damera S, Sarma S V., Inati SK, Zaghloul KA. Reinstatement of distributed cortical oscillations occurs with precise spatiotemporal dynamics during successful memory retrieval. Proc Natl Acad Sci. 2014;111: 18727–18732. doi:10.1073/pnas.1417017112

27. Maris E, Oostenveld R. Nonparametric statistical testing of EEG- and MEG-data. J Neurosci Methods. 2007;164: 177–190. doi:10.1016/j.jneumeth.2007.03.024

28. Näätänen R, Picton T. The N1 wave of the human electric and magnetic response to sound: A review and an analysis of the component structure. Psychophysiology. 1987;24: 375–425. doi:10.1111/j.1469-8986.1987.tb00311.x

29. Spiegler A, Hansen ECA, Bernard C, McIntosh AR, Jirsa VK. Selective activation of resting-state networks following focal stimulation in a connectome-based network model of the human brain. eNeuro. 2016;3. doi:10.1523/ENEURO.0068-16.2016

30. Lioumis P, Kičić D, Savolainen P, Mäkelä JP, Kähkönen S. Reproducibility of TMS-evoked EEG responses. Hum Brain Mapp. 2009;30: 1387–1396. doi:10.1002/hbm.20608

31. Kerwin LJ, Keller CJ, Wu W, Narayan M, Etkin A. Test-retest reliability of transcranial magnetic stimulation EEG evoked potentials. Brain Stimul. 2017; doi:10.1016/j.brs.2017.12.010

32. Soderlund GBW, Björk C, Gustafsson P. Comparing auditory noise treatment with stimulant medication on cognitive task performance in children with attention deficit hyperactivity disorder: Results from a pilot study. Front Psychol. 2016;7. doi:10.3389/fpsyg.2016.01331

33. Furnham A, Strbac L. Music is as distracting as noise: The differential distraction of background music and noise on the cognitive test performance of introverts and extraverts. Ergonomics. 2002;45: 203–217. doi:10.1080/00140130210121932

34. Herweg NA, Bunzeck N. Differential effects of white noise in cognitive and perceptual tasks. Front Psychol. 2015;6. doi:10.3389/fpsyg.2015.01639

35. Gordon PC, Zrenner C, Desideri D, Belardinelli P, Zrenner B, Brunoni AR, et al. Modulation of cortical responses by transcranial direct current stimulation of dorsolateral prefrontal cortex: A resting-state EEG and TMS-EEG study. Brain Stimul. 2018;11: 1024–1032. doi:10.1016/j.brs.2018.06.004

